# Molecular basis of SARS-CoV-2 Omicron variant receptor engagement and antibody evasion and neutralization

**DOI:** 10.1101/2022.01.10.475532

**Authors:** Qin Hong, Wenyu Han, Jiawei Li, Shiqi Xu, Yifan Wang, Zuyang Li, Yanxing Wang, Chao Zhang, Zhong Huang, Yao Cong

## Abstract

The SARS-CoV-2 Omicron variant exhibits striking immune evasion and is spreading globally at an unprecedented speed. Understanding the underlying structural basis of the high transmissibility and greatly enhanced immune evasion of Omicron is of high importance. Here through cryo-EM analysis, we present both the closed and open states of the Omicron spike, which appear more compact than the counterparts of the G614 strain, potentially related to the Omicron substitution induced enhanced protomer-protomer and S1-S2 interactions. The closed state showing dominant population may indicate a conformational masking mechanism of immune evasion for Omicron spike. Moreover, we capture two states for the Omicron S/ACE2 complex with S binding one or two ACE2s, revealing that the substitutions on the Omicron RBM result in new salt bridges/H-bonds and more favorable electrostatic surface properties, together strengthened interaction with ACE2, in line with the higher ACE2 affinity of the Omicron relative to the G614 strain. Furthermore, we determine cryo-EM structures of the Omicron S/S3H3 Fab, an antibody able to cross-neutralize major variants of concern including Omicron, elucidating the structural basis for S3H3-mediated broad-spectrum neutralization. Our findings shed new lights on the high transmissibility and immune evasion of the Omicron variant and may also inform design of broadly effective vaccines against emerging variants.

## Introduction

Severe acute respiratory syndrome coronavirus 2 (SARS-CoV-2) has undergone considerable evolution since its initial discovery in December 2019, leading to the emergence of a number of variants of concerns (VOCs) including Alpha (B.1.1.7)^1–5^, Beta (B.1.351)^4–8^, Gamma (P1)^9^, and Delta (B.1.617.2)^10,11^. These variants that harbor multiple mutations on their spike (S) protein show enhanced transmissibility and resistance to antibody neutralization^11^. Recently, a new variant, named Omicron (B.1.1.529), was first reported in South Africa in November 2021 and classified as the fifth VOC by the World Health Organization (WHO) on 26 November 2021. Omicron exhibits a high transmission rate (R0>3)^12,13^, and, as of 22 December 2021, it has spread into 110 countries^14^.

Omicron bears 37 mutations in its S protein relative to the original SARS-CoV-2 strain^15,16^. As the consequence, Omicron has been observed to extensively escape neutralization by previously developed neutralizing monoclonal antibodies (MAbs) or sera from vaccines or convalescent individuals^15,17–22^. Among all of the Omicron S mutations, 15 are present in the receptor-binding domain (RBD) that mediates the virus binding to its host-cell receptor— angiotensin-converting enzyme 2 (ACE2) and is also a major target for neutralizing antibodies^23–27^. In particular, 9 mutations are located within the receptor-binding motif (RBM) interacting directly with ACE2. However, Omicron still uses ACE2 as its entry receptor^22^. Moreover, the Omicron S appears to have an increased binding affinity to human ACE2 relative to the WT S^15,16,28^.

The high transmissibility and greatly enhanced resistance to antibody neutralization observed for Omicron makes this VOC particularly threatening. Therefore, further understanding of the nature of Omicron is of significant importance and may help in developing countermeasures against this VOC. The present study aimed to address from a structural aspect how Omicron binds the ACE2 receptor and how it recognizes or evades neutralizing antibodies raised against the original virus. We captured two cryo-EM structures of the Omicron S trimer in the closed and open state at 3.08- and 3.21-Å-resolution, respectively, revealing the Omicron spike is structurally more compact compared to the counterparts of the G614 strain. This could be related to the unique Omicron substitutions in SD1 and S2 regions. We also obtained two states of the Omicron S/ACE2 complex with S binding one or two ACE2s, respectively, suggesting that the substitutions on the RBM of Omicron result in formation of new salt bridges and H-bonds, as well as more complementary electrostatic surface properties. Moreover, we determined cryo-EM structures of the Omicron S in complex with the Fab of S3H3^29^, an antibody able to cross-neutralize major VOCs including Omicron, thus allowing elucidation of the structural basis for S3H3-mediated broad-spectrum neutralization.

## Results

### Closed and open state structures of the Omicron S trimer

To inspect the impact of the Omicron intense substitutions on the spike conformation, we prepared a prefusion-stabilized trimeric S protein of SARS-CoV-2 Omicron variant (Fig. S1) and subsequently determined its cryo-EM structures. Two cryo-EM maps, including an all RBD down conformation (termed Omicron S-close) and a one RBD-up open conformation (termed Omicron S-open), were obtained at 3.08- and 3.21-Å-resolution, respectively (Fig. 1A-B and Fig. S2A-C, Table S1). We then built an atomic model for each of the two structures (Fig. 1C, S2D). For the Omicron S-close state, the three protomers are well resolved and they display similar conformation with their RBDs in the down position (Fig. S2C and Fig. 1E). Strikingly, the Omicron S-close appears more twisted/compact in the RBDs relative to the G614 S-close structure (Fig. 1F)^30^. Also, in the Omicron S-open state, structural comparison showed that the RBDs are slightly more twisted/compact than that of the G614 S-open^30^ (Fig. 1G). There is no linoleic acid (termed LA) in the Omicron S-open and S-close maps, as in our recent Delta, Kappa, and Beta S structures obtained in the same construction and purification condition^31,32^. LA binding has been detected in the tightly closed WT S trimer structures^33–36^, and been suggested to lead to more compacted RBDs^33^. Collectively, the Omicron S trimer is more compact than that of G614, and this is not caused by LA binding. Moreover, in the Omicron S-open structure, the down RBD-3 is relative dynamic and less well resolved than RBD-2 (Fig. 1B). Our further 3D variability analysis (3DVA)^37^ on the Omicron S trimer dataset revealed an intrinsic rising up motion of RBD-1, which could alter the original RBD-1/-3 contact and destabilize RBD-3, making it extremely dynamic and may transiently rise up (Fig 1H, Movie S1).

**Fig. 1.**
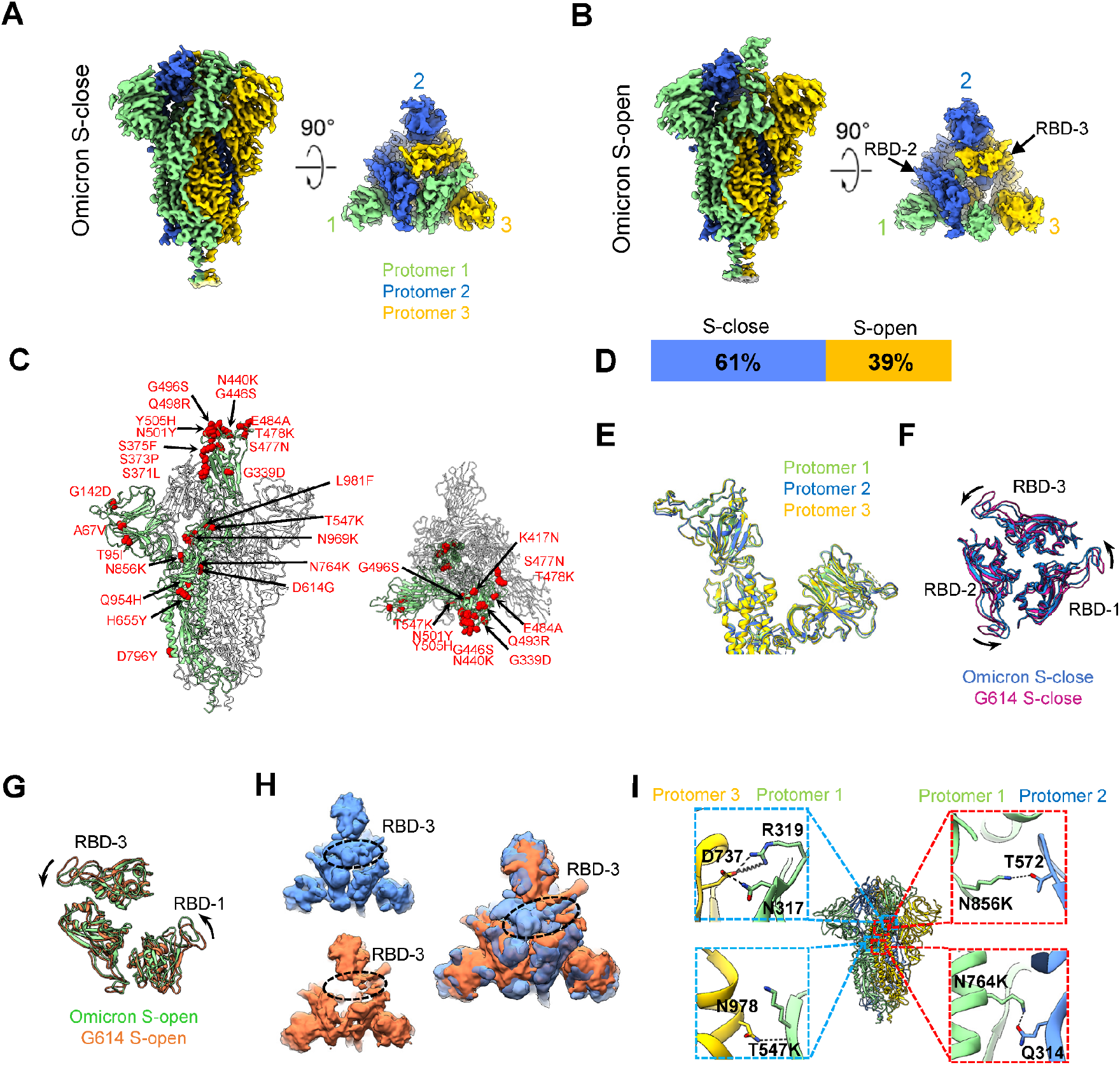
Cryo-EM structures of the SARS-CoV-2 Omicron S trimer. (A-B) Cryo-EM maps of the Omicron S-close (A) and S-open (B) state. Protomer 1, 2, and 3 are shown in light green, royal blue, and gold, respectively, which color scheme was followed throughout. (C) Atomic model of the Omicron S-open, with mutations indicated by red sphere and labeled. (D) Population distribution of the Omicron S-close and S-open. (E) Side view of the overlaid protomers of the Omicron S-close. (F) Top view of the overlaid RBDs of the Omicron S-close (blue) and the G614 S-close (PDB: 7KRQ, purple), indicating a twist of the Omicron S-close relative to that of G614. (G) Top view of the overlaid RBDs of the Omicron S-open (light green) and the G614 S-open (PDB: 7KRR, orange), indicating a twist of the Omicron S-open relative to that of G614. (H) One representative 3DVA motions of the Omicron S dataset. The left two maps illustrate the top view of two extremes in the motion with the RBD-3 indicated by dotted black ellipsoid, and the top view of the overlaid two extreme maps is shown in the right. (I) Newly formed H-bonds (black dashed line) and salt bridges (spring) in the interfaces of protomer 1/3 and protomer 1/2 of the S-close state.

Noteworthy, the population distribution of the Omicron S-close and S-open is about 61% and 39% (Fig. 1D), respectively, displaying a considerable population distribution shift to the closed state than that of the Kappa and Beta variants S trimer (both around 50%-50% open-transition ratio) or that of the Delta S (75.3%-24.7% open-transition ratio) from our recent studiues^31,32^. Taken together, the Omicron S trimer appears more prone to the closed state and potentially stabilized relative to the counterparts of the G614, Kappa, Beta, and Delta variants. To investigate the underlying molecular basis of this extra stability, we inspected the protomer interaction interface of Omicron S-close (Table S2, S3) and found three sets of new hydrogen bond (H-bond) and salt bridge interactions induced by the unique Omicron substitutions beyond the NTD/RBD regions (Fig. 1I). Specifically, the T547K from the SD1 of protomer 1 forms a new H-bond with the N978 from S2 of protomer 3, which could enhance the S1-S2 subunits interaction between the two protomers; the N856K and N764K from protomer 1 can form H-bonds with T572 and Q314 from protomer 2, respectively. We also observed multiple new H-bonds and salt bridges formed between the N317/R319 of protomer 1 and the D737 of protomer 3. These extra interactions mainly induced by Omicron substitutions in SD1 and S2 contribute greatly to the linkage/allosteric network between neighboring protomers and between the S1 and S2 subunits, markedly stabilizing the Omicron S trimer and inhibiting its transformation towards the fusion-prone open state and subsequent shielding of S1.

### Structural basis of enhanced S-ACE2 interaction for the Omicron variant

Compared with the WT strain, the Omicron variant bears 15 mutations in the RBD region, 9 of which are located in RBM^15^. We assessed whether these mutations affect the human ACE2 receptor-binding ability of the Omicron S trimer by performing biolayer interferometry (BLI) assay. The S trimers of the G614 and Delta variant were also analyzed for comparison purpose. We found that the ACE2-binding affinity of the Omicron S trimer (KD = 80 nM) is comparable to that of the Delta S (KD = 88 nM) but is about 3 folds higher than that of the G614 S (KD = 237 nM) (Fig. 2A), in consistence with the data from other recent preprints^15,16,28^.

**Fig. 2.**
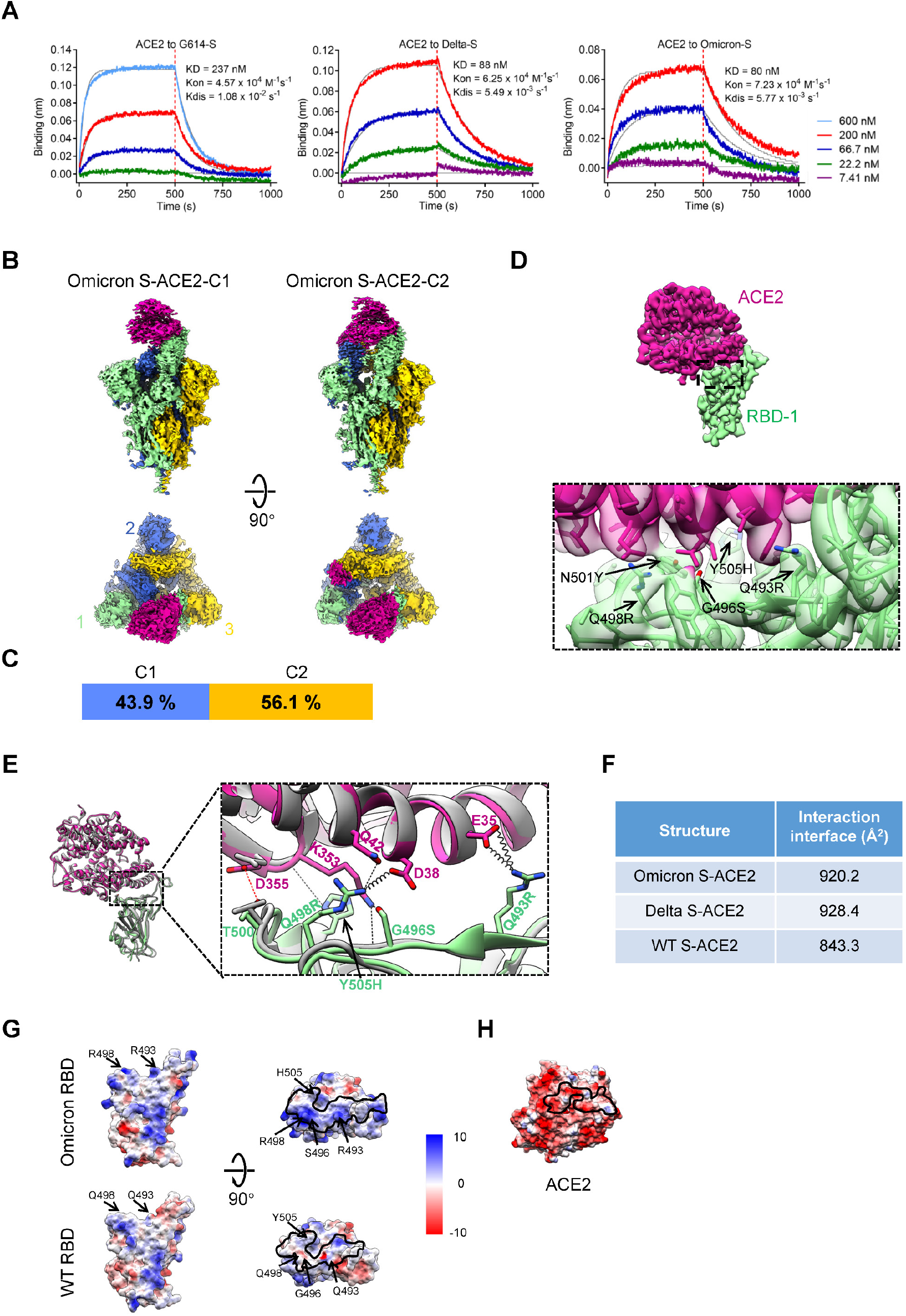
Structural basis of enhanced Omicron variant S trimer/ACE2 interaction. (A) Measurement of the binding affinity between ACE2 monomer and the S trimer of the G614 (left), Delta (middle) or Omicron (right) variants using bio-layer interferometry (BLI). Biotinylated S trimers were loaded onto streptavidin sensors and then allowed to interact with different concentrations of ACE2 (shown on the right). Raw sensorgrams and fitting curves were shown in color and gray, respectively. Association and dissociation phases were divided by red dotted lines. (B) Cryo-EM maps of the Omicron S-ACE2 complex in two distinct conformational states. ACE2 is shown in violet red. This color scheme is followed throughout. (C) Population distribution of the Omicron S-ACE2 conformers. (D) Density map of the focused refined Omicron RBD-1-ACE2 and the zoomed-in view of the RBD-ACE2 interaction interface, showing the side chain densities of the Q493R, G496S, Q498R, N501Y and Y505H on RBM. (E) The substituted residues R493, S496, R498 and H505 of Omicron RBM form new interactions with E35, D38, Q42 and K353 of ACE2 (spring represents salt bridge, and the black dashed line represents H-bond) relative to that in WT RBD-ACE2 (PDB: 6M0J, in dark grey). A newly formed H-bond without substitution is shown in red dashed line. (F) Interaction interface areas between ACE2 and RBD of Omicron, Delta (PDB: 7W9I), or WT (PDB: 6M0J), analyzed using PISA. (G) The electrostatic surface properties of Omicron and WT RBDs, with the mutated residues indicated. Black lines depict the footprint of ACE2 on RBD. (H) The electrostatic surface property of ACE2, with residues in proximity to RBD-1 (< 4 Å) indicated (Table. S5).

Next, we carried out cryo-EM study on the Omicron S trimer in complex with human ACE2 peptidase domain (PD) (Fig. S3). We obtained two cryo-EM maps (Fig. 2B), including a conformation with one RBD up and engaged with an ACE2 (termed Omicron S-ACE2-C1) and another one containing two “up” RBDs (RBD-1 and RBD-2) bound with ACE2 (termed Omicron S-ACE2-C2), at 3.69- and 3.66-Å-resolution, respectively (Fig. S4A-B, and Table S1). In the S-ACE2-C2 map, density of RBD-2-associated ACE2 appears weaker than that of the stably associated ACE2 on RBD-1. We then built an atomic model for each of the two structures (Fig. S4C). The population distribution between Omicron S-ACE2-C1 and -C2 is about 43.9% versus 56.1% (Fig. 2C), displaying an obvious higher one-RBD-up C1 population than that of the Beta/Kappa/Delta variants (C1 population ranges from 8.3% to 14.1%) observed in our recent studies^31,32^. These three variants also showed a C3 state with all-three-up RBDs associated with ACE2 (27.7% to 46.6% populated)^31,32^, not detected here in the Omicron variant, in line with recent preprint reports^16,28,38^. Taken together, the Omicron S trimer exhibits less ability to transform to the more RBD-up C2/C3 states as compared to that of the Beta, Kappa, and Delta VOCs.

To further understand the structural details of the RBD-ACE2 interaction interface, we focus-refined the stably associated Omicron RBD-1-ACE2 region to 3.67-Å-resolution (Fig. 2D and Fig. S4A). Inspection of this map revealed that many of the substitutions in RBM, including Q493R, G496S, Q498R, S477N, and Y505H, exhibit new interactions with ACE2 receptor compared with the interaction interface of the WT RBD-ACE2 (PDB: 6M0J)^26^. Specifically, Q493R with ACE2 E35 and Q498R with ACE2 D38 form three new salt bridges; G496S and Y505H both with ACE2 K353, Q498R with ACE2 Q42, and S477N with ACE2 Q19 form new H-bonds (Fig. 2D-E, Table. S4), generally in line with recent studies^16,28,38–41^. Moreover, we observed an extra H-bound between T500 and ACE2 D355 (Fig. 2E). Our previous research defined that Y505A obviously decreased ACE2 binding affinity^36^, so Y505H mutation in Omicron may maintain or even enhance ACE2 binding. In the meanwhile, the K417N substitution, occurred in Omicron as well as in Beta and Delta variants, is known to markedly reduce ACE2 binding through abolishing multiple salt bridges/H-bonds with ACE2 D30^26,42,43^. Together, these newly formed RBM-ACE2 interactions may compensate the loss of some original RBM-ACE2 interactions due to the residue changes introduced into the Omicron RBM.

Further inspection of the surface property showed that the substitutions in RBM, especially Q493R, G496S, Q498R, and Y505H, render the substituted site within the ACE2 interaction footprint more positively charged, which could strengthen the RBM interaction with the overall negatively charged ACE2 in the interaction interface (Fig. 2G-H). Corroborating this, the Omicron RBD-ACE2 interaction area (920.2 Å^2^) is enlarged compared to that of the WT (843.3 Å^2^), while it is comparable to that of the Delta RBD-ACE2 (928.4 Å^2^)^32^ (Fig. 2F). This is also in agreement with our BLI data showing that the ACE2-binding affinity of the Omicron S is similar to that of the Delta S but is higher than that of the G614 S (Fig. 2A).

### Sensitivity of Omicron to select neutralizing antibodies

We have generated a number of MAbs that potently neutralize the original SARS-CoV-2 strain in previous studies^29,44^. Five of these MAbs, including 2H2^44^, 3C1^44^, 8D3^44^, S5D2^29^, and S3H3^29^, were selected and tested in parallel for neutralization of the wild-type (WT, Wuhan-Hu-1 strain), Delta, or Omicron pseudoviruses. The neutralization data were shown in Fig. 3A-B. It was found that the IC50 values of MAbs 3C1, 2H2, 8D3, and S3H3 against Delta were comparable (less than 2.5-fold variation) to the corresponding ones against WT, whereas S5D2 was still neutralizing to Delta (IC50 = 734.6 ng/mL) but was about 90-fold less potent. In Omicron neutralization tests, three MAbs, 3C1, 8D3, and S5D2, lost neutralization activity (IC50 > 10 μg/mL). However, 2H2 and S3H3 remained highly effective against Omicron with IC50s being 30.4 and 53.3 ng/mL, respectively, despite that a 3.3-fold increase (relative to the WT) in IC50 value was observed for 2H2. These data demonstrate that 2H2 and S3H3 are two potent neutralizing MAbs against Omicron and also show that Omicron can more extensively escape antibody neutralization than Delta.

**Fig. 3.**
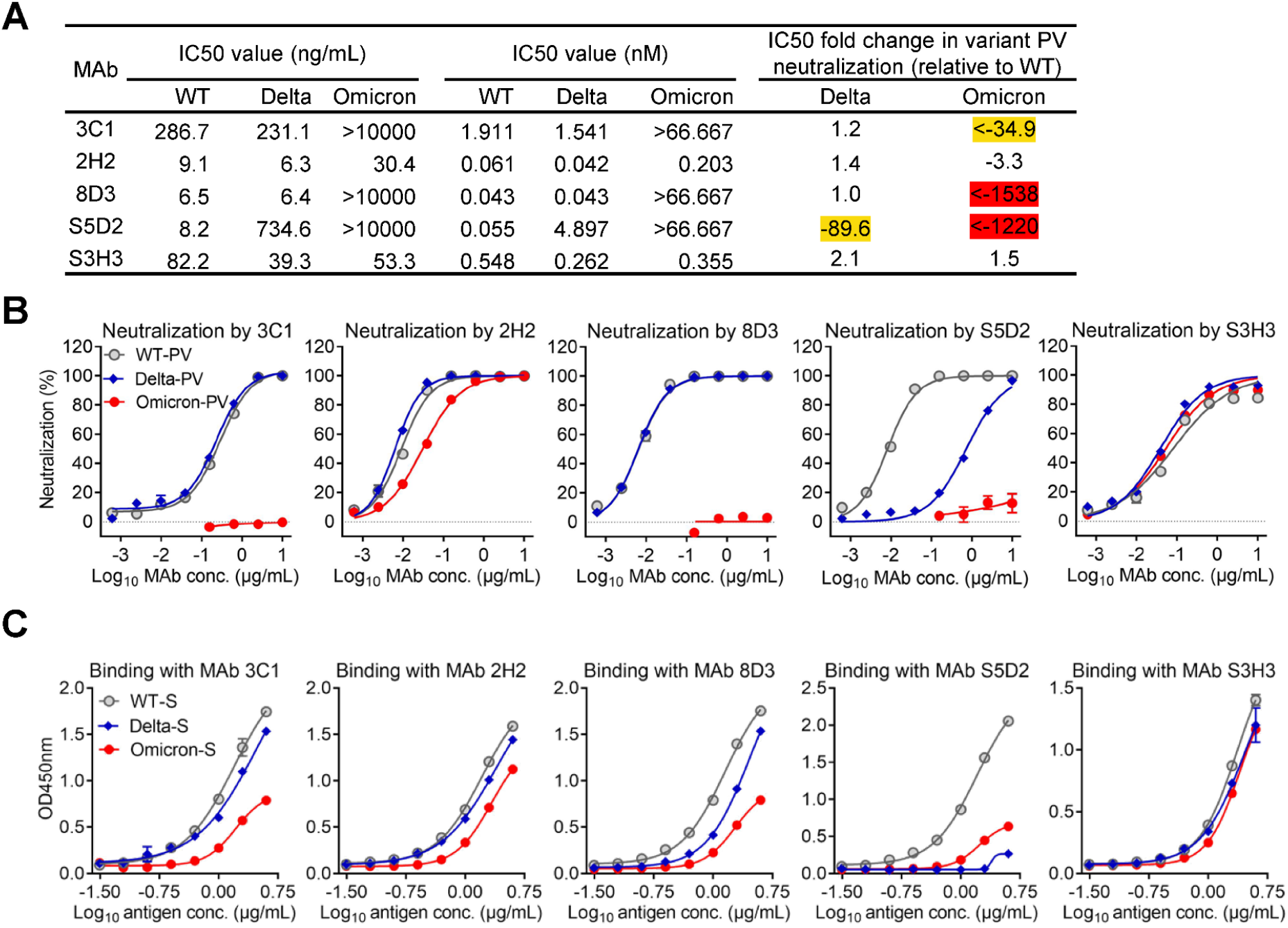
Neutralization and binding activities of the MAbs against SARS-CoV-2 Omicron and Delta variants. The MAbs were raised against WT RBD or S trimer proteins. (A) Neutralization IC50 values and fold changes in neutralization potency for Delta and Omicron variant pseudoviruses (PV) compared to WT pseudovirus. A minus sign (-) denotes decrease. Orange shade, more than 10-fold decrease; red shade, more than 1000-fold decrease. (B) Neutralization of the MAbs towards WT, Delta, and Omicron SARS-CoV-2 pseudoviruses. All MAbs were 4-fold serially diluted. Data are expressed as mean ± SEM of four replicate wells. (C) Binding activities of the MAbs to recombinant S trimers of the WT, Delta, and Omicron SARS-CoV-2 strains were tested by ELISA. Serially diluted S trimer proteins were coated onto the ELISA wells. Data are mean ± SD of triplicate wells.

We then compared the binding ability of the five MAbs to the WT, Delta, and Omicron S proteins by ELISA. As shown in Fig. 3C, for MAb S5D2, its binding to the Delta and to the Omicron S was nearly abolished; for MAbs 3C1 and 8D3, their reactivity profile with the Delta S closely resembled that towards the WT S but their binding to the Omicron S reduced significantly; for MAb 2H2, its binding curve to the Omicron S was similar to those towards the WT and Delta S despite the binding efficiency to the Omicron S was slightly lower; meanwhile, MAb S3H3 produced nearly identical binding curves to the three S proteins. Overall, the antigen-binding ability of the MAbs was in good agreement with their neutralization potency towards specific variant pseudovirus (Fig. 3A-C).

Collectively, the above results demonstrate that Omicron remains sensitive to binding and neutralization by MAbs 2H2 and S3H3 whereas it displays resistance to 3C1, 8D3, and S5D2.

### Structural basis of Omicron neutralization by a broadly neutralizing antibody S3H3

MAb S3H3 is a unique neutralizing antibody that binds the SD1 region of the WT S^29^. To understand the structural basis of Omicron neutralization by S3H3, we carried out cryo-EM study and obtained two structures of the SARS-CoV-2 Omicron S trimer in complex with S3H3 Fab in distinct conformational states (Fig. S5A). Both structures showed two engaged Fab densities on the SD1 region of protomer 2 and protomer 3, but with the RBD-1 in the up (termed Omicron S-open-S3H3) or down (termed Omicron S-close-S3H3) conformations (Fig. 4A-B). The two maps were resolved to the resolution of 3.48 Å and 3.64 Å, respectively (Fig. S5C-D and Table S1). We then built an atomic model for each of the two structures (Fig. S5B). Compared with the free Omicron S-open, the S trimer in the S-open-S3H3 structure exhibited a slight twist and the RBD-1 displayed a 9.1° downward rotation (Fig. 4C), making the S trimer seemingly less “open” as a whole. Meanwhile, the SD1 showed a slight downward rotation (Fig. 4C).

**Fig. 4.**
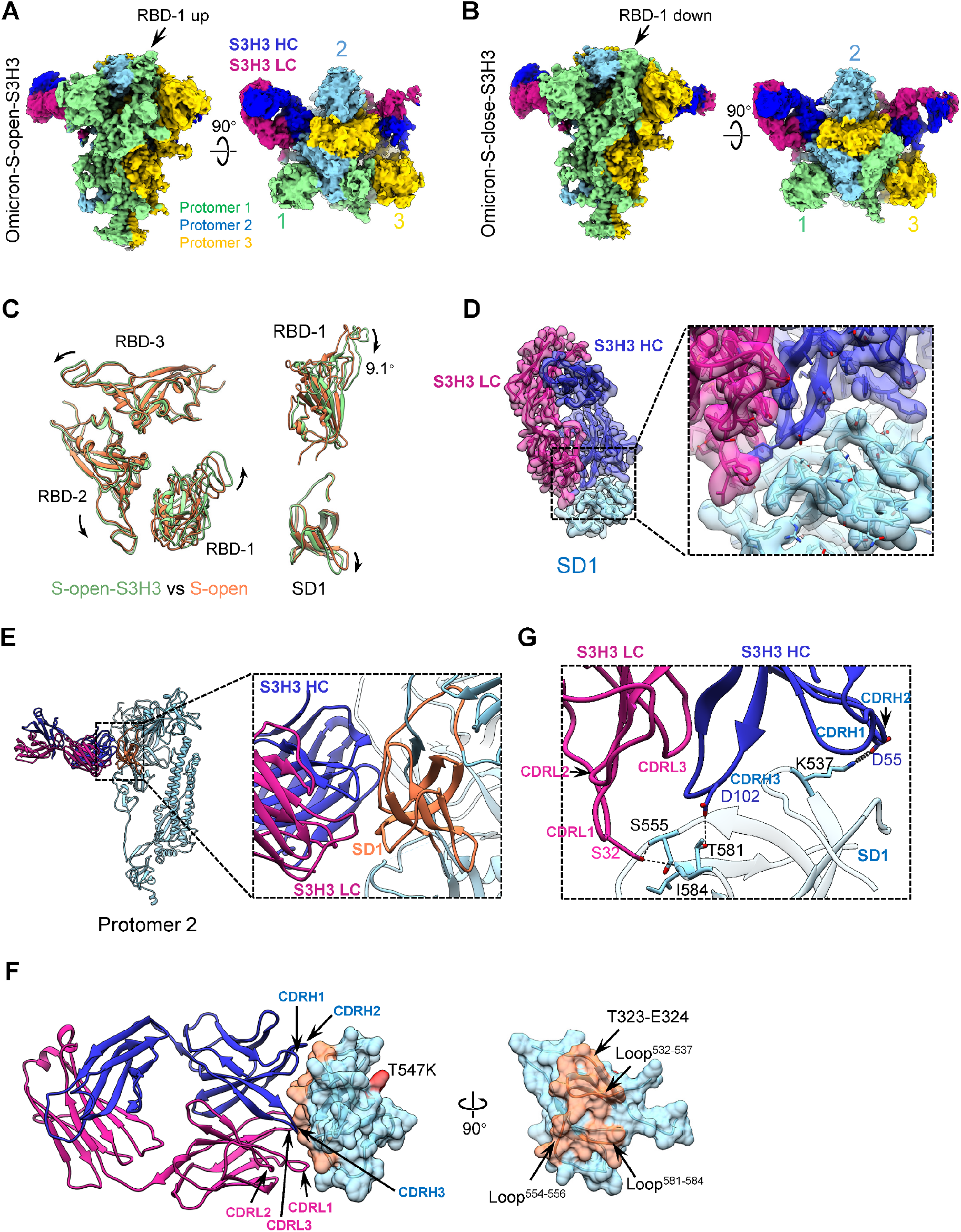
Cryo-EM analyses on the Omicron S-S3H3 Fab complex. (A-B) Side and top views of the cryo-EM map of the Omicron S-open-S3H3 (A) and S-close-S3H3 complex (B), with the heavy and light chains of S3H3 Fab in medium blue and violet red, respectively. The color scheme was followed. (C) Conformational comparison between Omicron S-open-S-S3H3 (light green) and Omicron S-open (orange), indicating a slight twist of the RBDs of S-open-S3H3 and the downward rotations of RBD-1 (up to 9.1°) and SD1. (D) Model-map fitting of the focus-refined Omicron SD1-S3H3 structure, and the zoomed-in view of the Omicron SD1-S3H3 interaction interface. The sidechain densities at the interface were well resolved. (E) The S3H3 binding on SD1 of protomer 2. (F) The interaction involved regions/residues between S3H3 Fab and SD1 with T547K labeled. (G) The SD1-S3H3 interaction interface analyzed using PISA, with major involved structural elements labeled (spring represents salt bridge, and the black dashed line represents H-bond).

To examine the interaction interface between the S3H3 Fab and the Omicron SD1, we further focused refined the SD1-S3H3 Fab region and obtained a map at 3.61-Å-resolution, with most of the sidechain densities well resolved (Fig. 4D). Our structural analysis suggested that the heavy chain of S3H3 Fab contributes more to the interactions with SD1 than the light chain does, i.e., all the three heavy-chain CDRs of S3H3 and its CDRL1 and CDRL3 interact with T323-E324 and the three loops (loop^532-537^, loop^554-556^, and loop^581-584^) of SD1 (Fig. 4E-F and Table. S7). Specifically, the S32 of CDRL1 forms H-bonds with the S555 and I584 of SD1, respectively, the D102 of CDRH3 forms a H-bond with the T581 from loop^581-584^, and the D55 of CDRH2 forms a salt bridge with the K537 from loop^532-537^ (Fig. 4G and Table. S6), thus constituting an intense interaction network between S3H3 Fab and SD1. A single mutation, T547K, is present in the SD1 region of Omicron, however, this mutation locates outside the footprint of S3H3 (Fig. 4F), thus will not affect the interaction between S3H3 and Omicron S. Collectively, S3H3 binds the extremely conserved SD1 region, therefore retains binding and neutralizing activity towards major VOCs including Omicron.

## Discussion

The SARS-CoV-2 Omicron variant has replaced the Delta variant and is now the predominant circulating VOC in many countries^14^. With 37 mutations in its spike, this variant shows striking immune evasion while it also displays increased binding affinity with human ACE2 relative to the WT strain^15,16,28^. Recent reports also showed that the Omicron S exhibits reduced furin cleavage and less S1 shedding^45,46^. It is essential to understand how the mutations present in the Omicron S contribute to the higher transmissibility and immune escape observed for this threatening variant. In this study, we performed cryo-EM study and biochemical analysis on the Omicron S trimer and its complex with ACE2 receptor or a broadly neutralizing antibody S3H3. We captured both the closed and the open states of the Omicron S trimer (Fig. 1A-B). In contrast to the S trimer of Delta/Beta/Kappa variants^31,32^, the Omicron S-close and S-open structures appear more twisted/compact than their counterpart of the G614 strain (Fig. 1F-G). This could be related to the unique Omicron substitution (T547K, N856K, and N764K in SD1 and S2)–induced enhanced interactions between neighboring protomers and between S1 and S2 subunits (Fig. 1I), which may hinder its spike transformation towards the fusion-prone open state and shielding of S1.

Noteworthy, our cryo-EM analysis revealed the dominantly populated (61%) conformation for the Omicron S trimer is in the closed state with all the RBDs buried, resulting in conformational masking preventing antibody binding and neutralization at sites of receptor binding, similar to that described for HIV-1 envelope^47,48^. This Omicron conformational masking mechanism of neutralization escape could affect all antibodies that bind to the up RBDs (such as class 1, 2, and 4 RBD antibodies^49^). While for Delta S trimer, our recent work showed the open-transition ratio is 75.3%-24.7%, indicating the conformational masking mechanism may be less effective for the Delta variant^32,36^. This could contribute greatly to the striking immune evasion of the Omicron variant^15,17–22^.

We then captured two states for the Omicron S/ACE2 complex with S binding one or two ACE2s under our experimental conditions (Fig. 2B-C). However, unlike the Delta S which tends to bind three ACE2 in majority^32^, Omicron binds up to two ACE2s. Further focus-refined RBD-1/ACE2 structure demonstrated that the substitutions on the RBM of Omicron (especially Q493R, G496S, Q498R, S477N, and Y505H) result in formation of new salt bridges and H-bonds, as well as more complementary electrostatic surface properties (Fig. 2E-H), which together may compensate abolished original RBM-ACE2 interactions^26,42,43^, leading to enhanced interactions with ACE2 and potentially enhanced transmissibility of the Omicron variant.

SARS-CoV-2 variants gain series of mutations in their S proteins, including RBD and NTD. As a consequence, VOCs significantly impact the potency of neutralizing antibodies originally developed against WT strains^50,51^. Omicron contains specific alterations that have previously been shown to impact vaccine resistance and also some newly introduced mutations. It is thus important to determine whether antibodies capable of neutralizing Omicron exist and if yes where they target. In the present study, we screened a panel of five previously isolated and characterized neutralizing MAbs^44^ for their potency against Omicron and Delta variants. Our results show that 2H2 and S3H3 retain potent neutralization towards Omicron and Delta (Fig. 3). Further structural study on the Omicron S-S3H3 Fab complex revealed a unique binding epitope of S3H3 within the SD1 region which links the S1 and S2 domains (Fig. 4A-E). S3H3 binding to S trimer may function as a “lock” to block the releasing of S1 from S2, resulting in inhibition of virus entry. The SD1 region targeted by S3H3 is extremely conserved among SARS-CoV-2 strains, with only one mutation T547K (which is away from the S3H3 binding footprint) present in Omicron (Fig. 4F), thus explaining the cross-neutralization ability of S3H3 towards Omicron, Delta, and other variants^29^. These findings also suggest a possibility to design SD1-based broad-spectrum SARS-CoV-2 vaccines.

In summary, the present study reveals that the Omicron spike is structurally more compact compared to the counterparts of other VOCs and has the likelihood to associate with fewer ACE2. The compact S-close state with dominant population may indicate a conformational masking mechanism of immune evasion for Omicron spike. However, the Omicron spike still maintains strong affinity to ACE2 due to an increased RBM-ACE2 interaction network contributed by new H-bonds/salt bridges and more favorable surface properties, thus providing a possible explanation to the high transmissibility of Omicron. In addition, our work shows that Omicron is able to escape majority of the RBD-directed MAbs owing to a relatively large number of residue changes in RBD and conformational masking, however, this variant remains sensitive to the SD1-targeting neutralizing MAb S3H3. Our findings provide structural insights into how Omicron maintains high transmissibility while greatly evades immunity, and may also inform design of broadly effective vaccines against emerging variants.

## Method

### Expression and purification of recombinant proteins

To express SARS-CoV-2 Omicron variant S glycoprotein ectodomain, the mammalian codon-optimized gene coding SARS-CoV-2 (hCoV-19 Botswana R42B90_BHP_000842207 2021, GISAID ID: EPI_ISL_6752027) S glycoprotein ectodomain (residues M1-Q1208) with proline substitutions at K986 and V987, a “GSAS” substitution at the furin cleavage site (R682–R685) was cloned into vector pcDNA 3.1+. A C-terminal T4 fibritin trimerization motif, a TEV protease cleavage site, a FLAG tag and a His tag were cloned downstream of the S glycoprotein ectodomain (Fig. S1A). The constructs of prefusion-stabilized S proteins of SARS-CoV-2 G614 and Delta (B.1.617.2) variants were prepared as previously reported^32^. A gene encoding human ACE2 PD domain (Q18-D615) with an N-terminal interleukin-10 (IL-10) signal peptide and a C-terminal His tag was cloned into vector pcDNA 3.4^36^. The recombinant proteins were prepared as the published protocol^36^. Briefly, the constructs were transiently transfected into HEK293F cells using polyethylenimine (PEI). Three days after transfection, the supernatants were harvested by centrifugation, and then passed through 0.45 μm filter membrane. The clarified supernatants were added with 20 mM Tris-HCl pH 7.5, 200 mM NaCl, 20 mM imidazole, 4 mM MgCl_2_, and incubated with Ni-NTA resin at 4°C for 1 hour. The Ni-NTA resin was recovered and washed with 20 mM Tris-HCl pH 7.5, 200 mM NaCl, 20 mM imidazole. The protein was eluted by 20 mM Tris-HCl pH 7.5, 200 mM NaCl, 250 mM imidazole.

### Bio-layer interferometry (BLI) assay

Before BLI assay, Ni-NTA purified recombinant S trimer proteins of the G614, Delta and Omicron SARS-CoV-2 variants were further purified by gel filtration chromatography using a Superose 6 increase 10/300 GL column (GE Healthcare) pre-equilibrated with PBS. Then, the S trimer proteins were biotinylated using the EZ-Link™ Sulfo-NHS-LC-LC-Biotin kit (Thermo Fisher) and then purified by Zeba™ spin desalting columns (Thermo Fisher).

Binding affinities of S trimers to ACE2 were determined by BLI analysis on an Octet Red96 instrument (Pall FortéBio, USA). Briefly, biotinylated S trimer proteins were immobilized onto streptavidin (SA) biosensors (Pall FortéBio). After washing with kinetic buffer (0.01 M PBS with 0.02% Tween 20 and 0.1% bovine serum albumin), these sensors were incubated with 3-fold serial dilutions of ACE2 monomer protein for 500 s. Subsequently, the biosensors were allowed to dissociate in kinetic buffer for 500 s. The data were analyzed using the Octet Data Analysis 11.0 software to calculate affinity constants.

### Neutralization

Luciferase (Luc)-expressing pseudoviruses bearing SARS-CoV-2 S proteins were constructed based on the HIV-1 backbone. Briefly, HEK 293T cells in 10-cm dish were co-transfected using PEI (polysciences) with 10 μg of Pcmv-Dr8.91 packaging plasmid, 10 μg of recombinant Plvx-IRES-ZsGreen1 plasmid containing luciferase reporter gene, and 2 μg of recombinant Pvax1 plasmids encoding SARS-CoV-2 S proteins. The cells were incubated with the transfection mixture for 6 h, and then 5 mL of fresh DMEM medium with 10% FBS was added to each dish. After incubation overnight, the media in the dishes was replaced with fresh DMEM medium (10% FBS). At 48 h post-transfection, the culture supernatant was harvested and frozen at −80 °C before use.

All MAbs were 4-fold serially diluted and tested by pseudovirus neutralization assay with human ACE2-overexpressing HEK 293T cells (293T-Hace2) following our previous protocol^44^. Two days after pseudovirus infection, luciferase activity was measured. Data were analyzed by non-linear regression using GraphPad Prism 8 to calculate half inhibitory concentration (IC50).

### ELISA

To test binding activities of recombinant Omicron S protein with our previously developed anti-SARS-CoV-2 MAbs^29,44^, recombinant S trimer proteins from WT^44^, Delta, or Omicron SARS-CoV-2 strains were 2-fold serially diluted and coated onto ELISA plates at 37 °C for 2 h. The plates were blocked with 5% milk in PBS-Tween 20 (PBST) at 37 °C for 1 h. After washing with PBST, the plates were incubated with 50 ng/well of each of the anti-SARS-CoV-2 MAbs^29,44^ at 37 °C for 2 h, followed by horseradish peroxidase (HRP)-conjugated anti-mouse IgG (Sigma, 1/5,000 dilution) at 37 °C for 1 h. After washing and color development, absorbance was measured at 450 nm. ELISA data were analyzed by non-linear regression using GraphPad Prism 8.

### Omicron S trimer/S3H3 Fab complex formation

The Omicron variant S trimer/S3H3 Fab complex was prepared following our previously reported protocol^29^. Briefly, purified S3H3 IgG was incubated with papain (300:1 W/W) in PBS buffer (in the presence of 20 mM L-cysteine and 1 mM EDTA) for 3 h at 37°C. The reaction was quenched by 20 mM iodoacetamide. Fab was purified by running over a HiTrap DEAE FF column (GE Healthcare) pre-equilibrated with PBS. Omicron S protein was incubated with S3H3 Fab in a 1:6 molar ratio on ice for 1 h. The Omicron S-S3H3 Fab complex was purified by size-exclusion chromatography using Superose 6 increase 10/300 GL column (GE Healthcare) in 20 mM Tris-HCl pH 7.5, 200 mM NaCl, 4% glycerol. The complex peak fractions were concentrated and assessed by SDS-PAGE.

### Cryo-EM sample preparation

To prepare the cryo-EM sample of the Omicron S trimer, a 2.2 μl aliquot of the sample was applied on a plasma-cleaned holey carbon grid (R 1.2/1.3, Cu, 200 mesh; Quantifoil). The grid was blotted with Vitrobot Mark IV (Thermo Fisher Scientific) at 100% humidity and 8 °C, and then plunged into liquid ethane cooled by liquid nitrogen. To prepare the cryo-EM sample of the Omicron S-ACE2 complex, purified Omicron S trimer was incubated with ACE2 in a 1:4 molar ratio on ice for 20 min and then vitrified using the same condition. The purified Omicron S-S3H3 complex was vitrified using the same procedure as for the Omicron S sample.

### Cryo-EM data collection

Cryo-EM movies of the samples were collected on a Titan Krios electron microscope (Thermo Fisher Scientific) operated at an accelerating voltage of 300 kV. For the three datasets, the movies were collected at a magnification of 64,000× and recorded on a K3 direct electron detector (Gatan) operated in the counting mode (yielding a pixel size of 1.093 Å), and under a low-dose condition in an automatic manner using EPU software (Thermo Fisher Scientific). Each frame was exposed for 0.1 s, and the total accumulation time was 3 s, leading to a total accumulated dose of 50.2 e^−^/Å^2^ on the specimen.

### Cryo-EM 3D reconstruction

For each dataset, the motion correction of image stack was performed using the embedded module of Motioncor2 in Relion 3.1^36,52,53^ and CTF parameters were determined using CTFFIND4^54^ before further data processing. Unless otherwise described, the data processing was performed in Relion3.1.

For the Omicron S dataset (Fig. S2), 600,845 particles remained after reference-free 2D classification in cryoSPARC v3.3.1^37^. After 3D classification and focused 3D classification on the RBD-1 region, we obtained an Omicron S-close map from 69,873 particles and an S-open map from 108,509 particles. After Bayesian polishing and CTF refinement, the Omicron S-open and S-close datasets were independently loaded into cryoSPARC v3.3.1^37^ and refined to the resolutions of 3.21 Å and 3.08 Å, respectively, using Non-uniform refinement. The overall resolution was determined based on the gold-standard criterion using a Fourier shell correlation (FSC) of 0.143. Moreover, we performed 3D Variability analysis (3DVA) on the Omicron S trimer dataset in cryoSPARC to capture its continuous conformational dynamics^37^.

For the Omicron S-ACE2 dataset (Fig. S3), 1,268,072 particles remained after reference-free 2D classification. After two rounds of 3D classification and further focused 3D classification on the RBD-1-ACE2 region, we obtained an Omicron S-ACE2 map from 141,538 particles. After Bayesian polishing and CTF refinement, the map was reconstructed to 3.53-Å-resolution. We then focused on RBD-2 for further classification and obtained two conformations with RBD-2 in the “down” or “up” position, termed S-ACE2-C1 and -C2, respectively. The two datasets were independently loaded into cryoSPARC v3.3.1 and refined using Non-uniform refinement to 3.69- and 3.66-Å-resolution, respectively. The overall resolution was determined based on the gold-standard criterion using a Fourier shell correlation (FSC) of 0.143. Here, after obtaining the 3.53-Å-resolution map of Omicron S-ACE2, we performed further local refinement on the RBD-1-ACE2 region in cryoSPARC to acquire a 3.67-Å-resolution map of this region.

For the Omicron S-S3H3 dataset (Fig. S5), similar data processing procedure was adapted as described for the Omicron S dataset to obtain a 3.5-Å-resolution S-S3H3 map from 238,162 particles. We then carried out focused 3D classification on the RBD-1 region, followed by Non-uniform refinement in cryoSPARC, and obtained a 3.48-Å-resolution S-open-S3H3 map from 162,221 particles and a 3.64-Å-resolution S-close-S3H3 map from 75,900 particles. In addition, after obtaining the 3.5-Å-resolution map, we performed focused 3D classification on the S3H3-SD1 region of protomer 2 (highlighted by dotted orange ellipsoid), leading to a dataset of 101,192 particles, which was further local refined on the S3H3-SD1 region in cryoSPARC, deducing a 3.61-Å-resolution map of this region. All of the obtained maps were post-processed through deepEMhancer^62^.

### Atomic model building

To build an atomic model for the Omicron S-open structure, we used the atomic model of Delta S-open (PDB: 7W92) from our prior study as the initial model^32^. We first fit the model into our Omicron S-open map in Chimera by rigid body fitting, then manually substituted the mutations of the Omicron variant in COOT^55^. Subsequently, we flexibly refined the model against our Omicron S-open map using ROSETTA^56^. Finally, we used the phenix.real_space_refine module in Phenix for the S trimer model refinement against the map^57^. For the S-close model, we utilized the down protomer from our recent Delta S-transition (PDB: 7W94)^32^ structure as initial template, and followed similar procedure described above for model refinement. For the Omicron S-ACE2 and the local refined RBD-1-ACE2 structures, we used the Delta S-ACE2 model (PDB: 7W98, 7W9I)^32^ as initial template, and followed similar procedure described above for model refinement. For the Omicron S-S3H3 and the local refined RBD-1-S3H3 structures, we utilized our recent Beta S-S3H3 model (PDB: 7WDF)^29^ as template, and followed similar procedure described above for model refinement. The atomic models were validated using Phenix.molprobity command in Phenix. Interaction interface analyses were conducted through PISA server^58^.

UCSF Chimera and ChimeraX were applied for figure generation, rotation measurement, and coulombic potential surface analysis^59,60^.

## Acknowledgements

We are grateful to the staffs of the NCPSS Electron Microscopy facility, Database and Computing facility, and Protein Expression and Purification facility for instrument support and technical assistance. This work was supported by grants from the Strategic Priority Research Program of CAS (XDB37040103 and XDB29040300), National Key R&D Program of China (2017YFA0503503 and 2020YFC0845900), the NSFC (32130056 and 31872714), the NSFC-ISF 31861143028, Shanghai Academic Research Leader (20XD1404200). Chao Zhang is supported by the Youth Innovation Promotion Association of the Chinese Academy of Sciences (CAS) and Shanghai Rising-Star Program (21QA1410000).

## Author contributions

Y.C. and Z.H. designed the experiments; Y-X. Wang expressed and purified the proteins with assistants of Z.L. and S.X.; Q.H. and W.H. performed cryo-EM data acquisitions; Q.H., W.H., J.L., and Y-F. Wang performed cryo-EM reconstructions, model buildings; C.Z. and S.X. performed biochemical analyses; J.L. Q.H., W.H., Y-F. Wang and C.Z. analyzed the data; Y.C. and Z.H. together with Q.H., J.L. W.H., Y-X. Wang and C.Z. wrote the manuscript.

## Data availability

All data presented in this study are available within the figures and in the Supplementary Information. Cryo-EM maps determined for the SARS-CoV-2 Omicron S trimer have been deposited at the Electron Microscopy Data Bank with accession codes EMD-32556 and EMD-32557, and the associated atomic models have been deposited in the Protein Data Bank with accession codes 7WK2 and 7WK3 for S-open and S-close, respectively. For the S-ACE2 dataset, related cryo-EM maps have been deposited in the Electron Microscopy Data Bank with accession codes EMD-32558, EMD-32559 and EMD-32560, and the associated models have been deposited in the Protein Data Bank with accession codes 7WK4, 7WK5 and 7WK6 for S-ACE2-C1, S-ACE2-C2 and RBD-1-ACE2, respectively. For the S-S3H3 Fab dataset, related cryo-EM maps have been deposited in the Electron Microscopy Data Bank with accession codes EMD-32562, EMD-32563, and EMD-32564, and the associated models have been deposited in the Protein Data Bank with accession codes 7WK8, 7WK9 and 7WKA for SD1-S3H3, S-open-S3H3 and S-close-S3H3, respectively.

## Supplemental figures

**Fig. S1.**
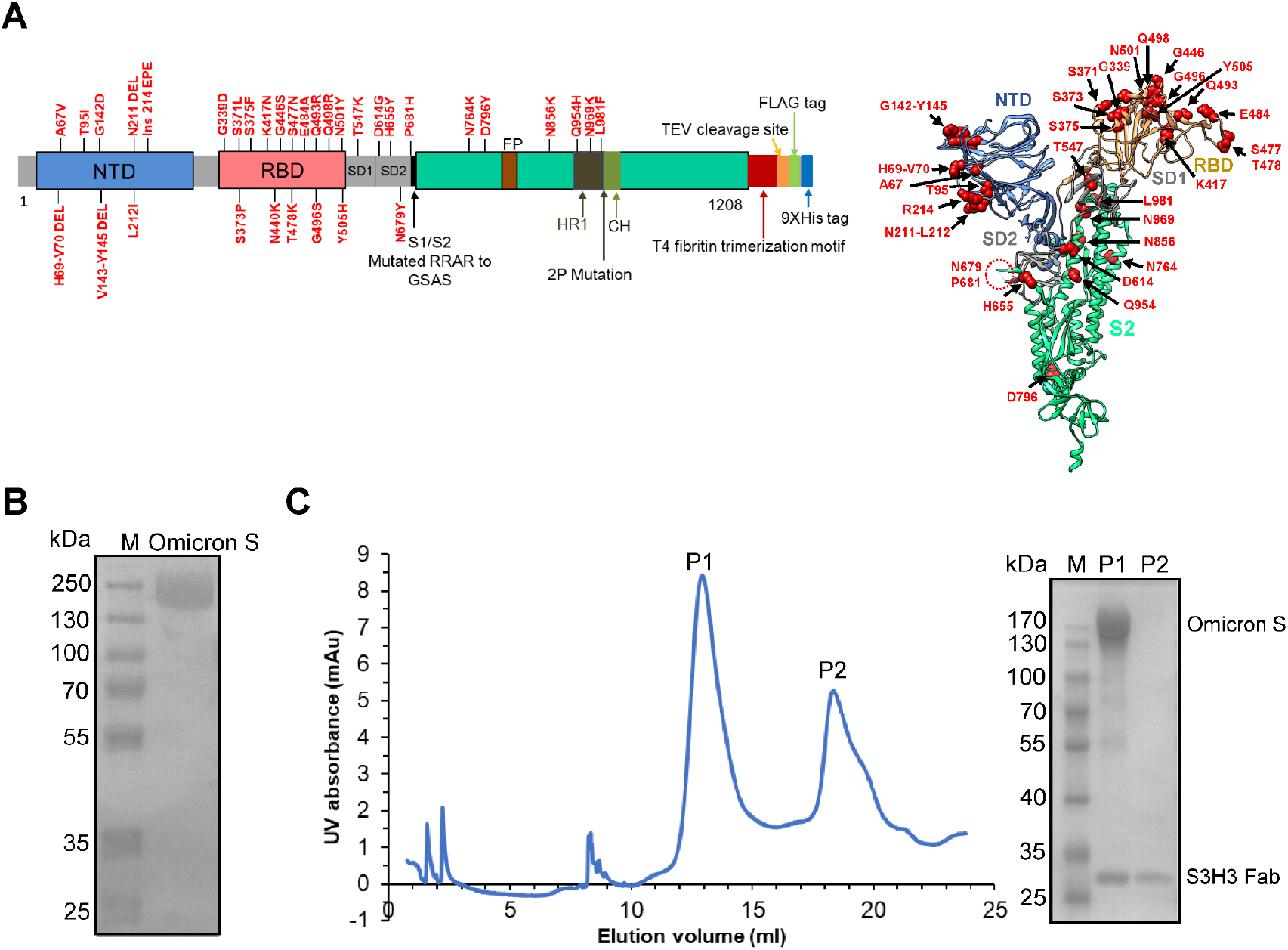
Purification of Omicron variant S and S-S3H3 Fab complex. (A) Schematic diagram of the Omicron variant S organization in this study (left, positions of all mutations are indicated), and the model of a SARS-CoV-2 S protomer (right) with mutation sites of the Omicron variant shown as red sphere. (B) SDS-PAGE analysis of the purified Omicron variant S protein. (C) Size-exclusion chromatogram and SDS-PAGE analysis of the formed Omicron S-S3H3 Fab complex.

**Fig. S2.**
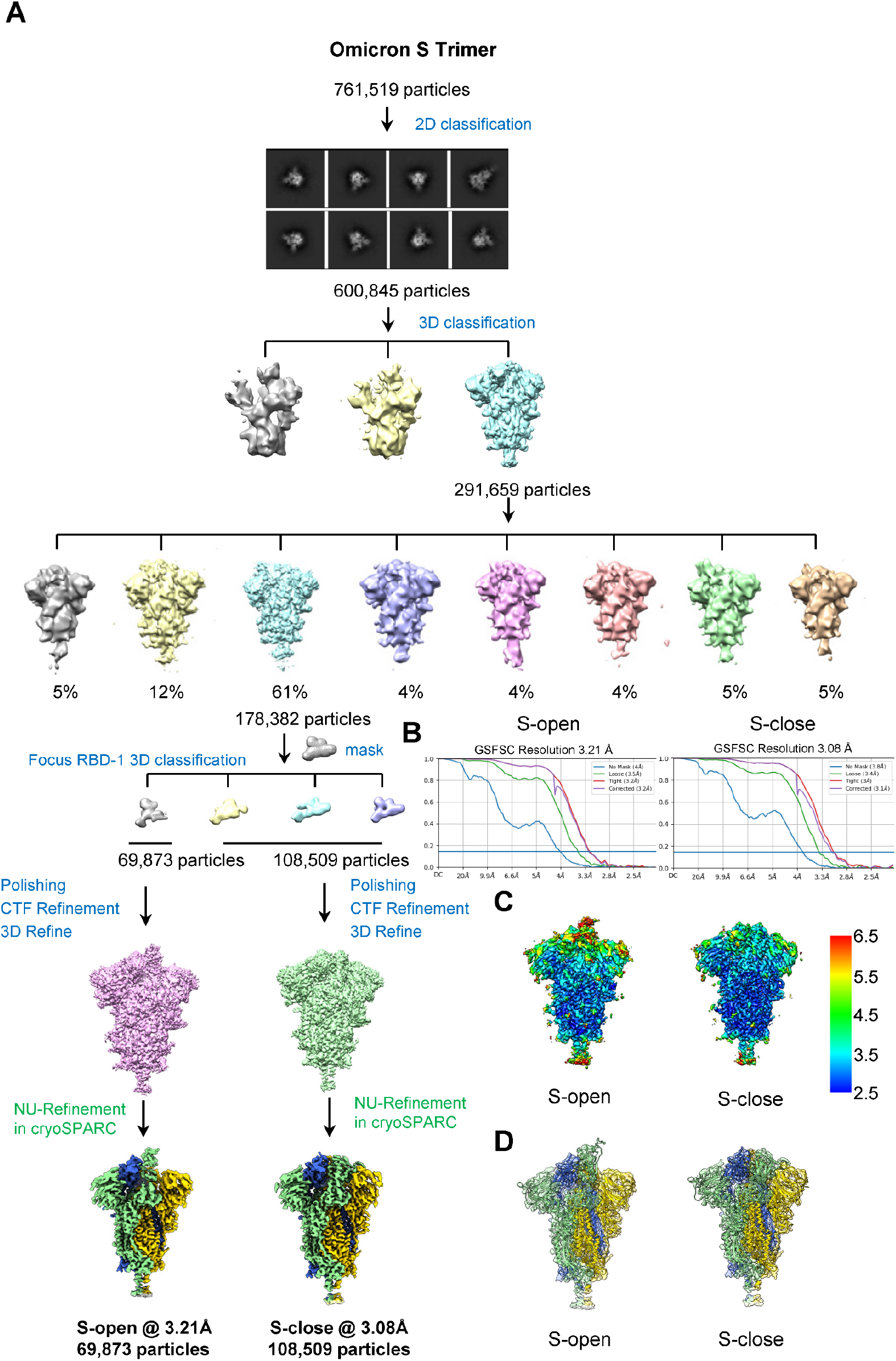
Cryo-EM analysis on the Omicron S trimer. (A) Data processing workflow for structure determination of the Omicron S trimer. The reference-free 2D class averages are also presented. (B) Resolution assessment of Omicron S-open and S-close maps by FSC at 0.143 criterion. (C) Local resolution evaluation of the Omicron S-open and S-close maps. (D) Model-map fitting of the Omicron S-open and S-close structures.

**Fig. S3.**
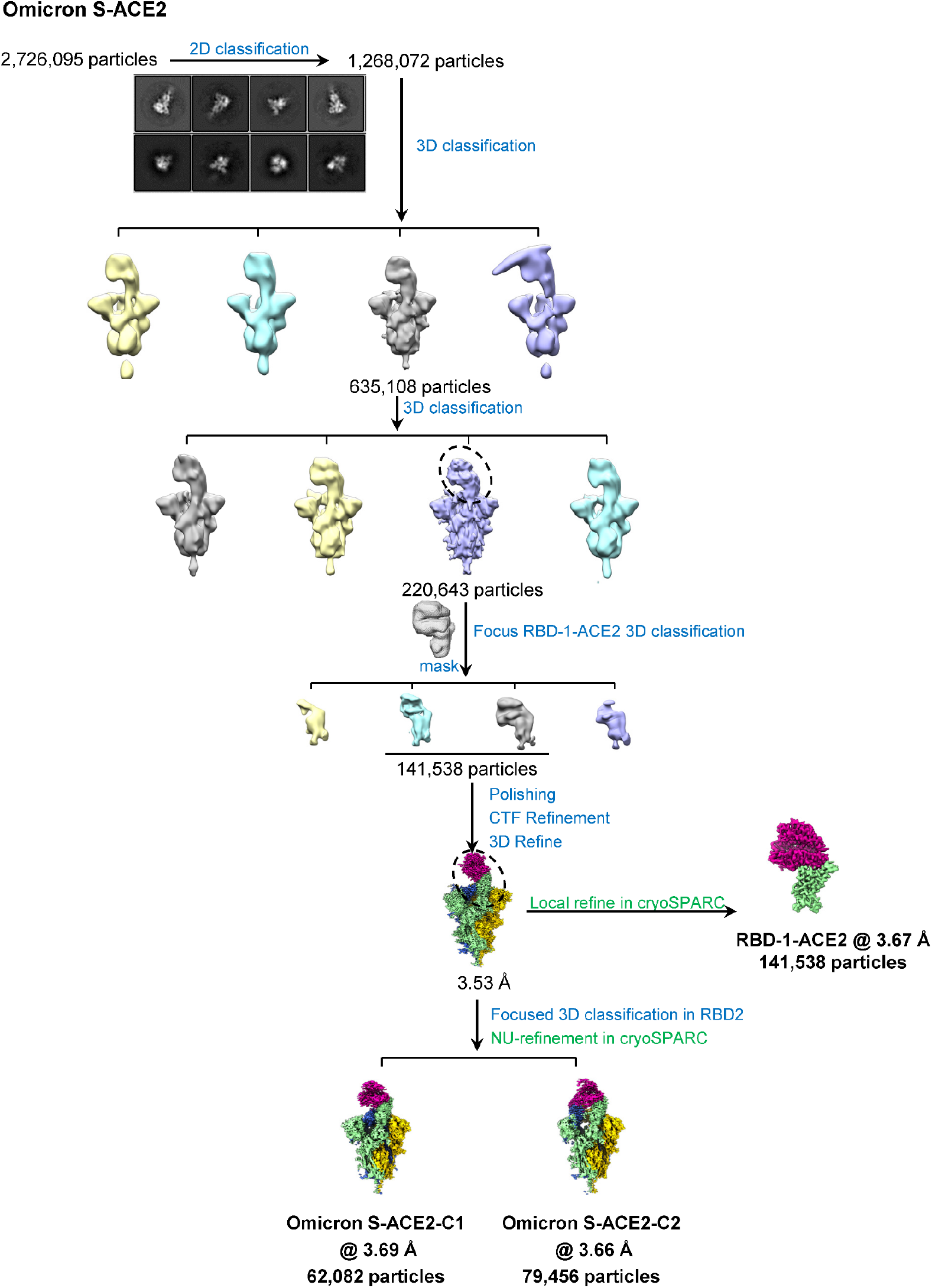
Cryo-EM data processing procedure for the Omicron S-ACE2 complex.

**Fig. S4.**
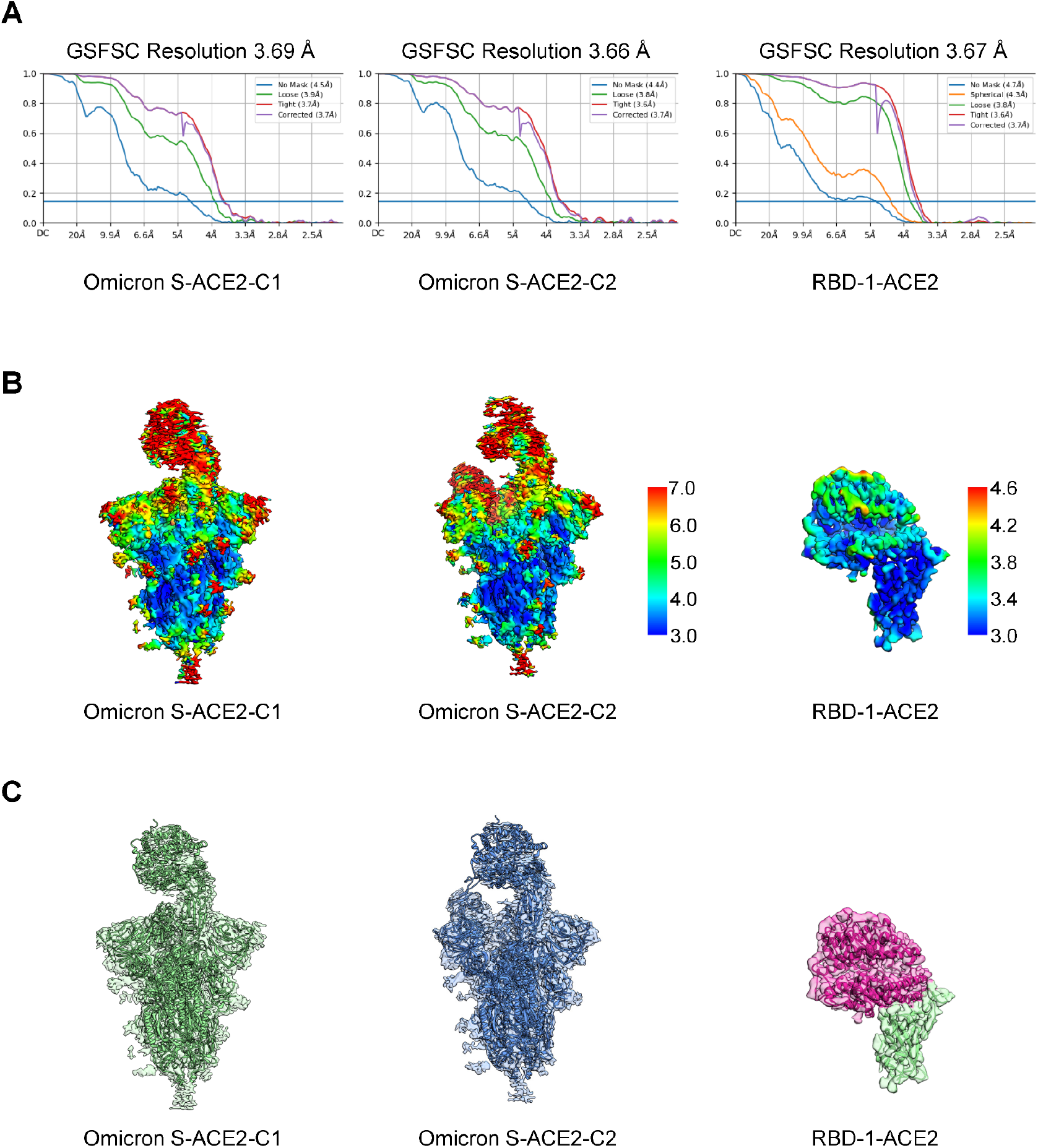
Cryo-EM analysis on the Omicron S-ACE2 complex. (A) Resolution assessment of the cryo-EM maps by FSC at 0.143 criterion. (B-C) Local resolution evaluation (B) and Model-map fitting (C) for the Omicron S-ACE2 complex maps and the RBD-1-ACE2 map.

**Fig. S5.**
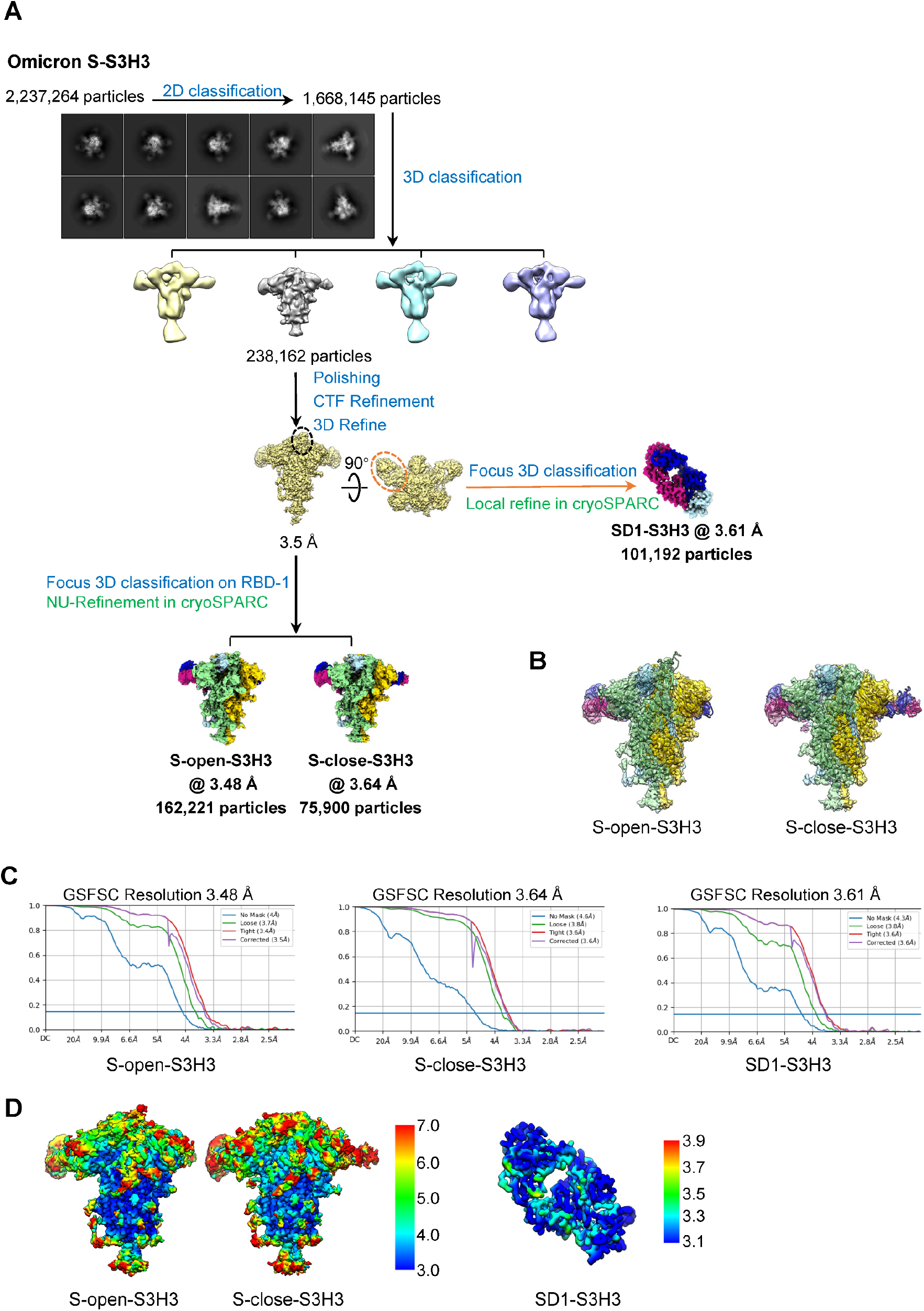
Cryo-EM analysis on the Omicron S-S3H3 Fab complex. (A) Data processing workflow for the Omicron S-S3H3 Fab complex. The reference-free 2D class averages are also presented. (B) Model-map fitting for the Omicron S-S3H3 complex. (C) Resolution assessment of the cryo-EM maps by FSC at 0.143 criterion. (D) Local resolution evaluation of the Omicron S-S3H3 and SD1-S3H3 maps.

## Supplemental tables

**Table S1.** Cryo-EM data collection and refinement statistics for Omicron S, Omicron S-ACE2, and Omicron S-S3H3.

|  |  |  |  |  |  |  |  |  |
| --- | --- | --- | --- | --- | --- | --- | --- | --- |
|  | Omicron S |  | Omicron S-ACE2 |  |  | Omicron S-S3H3 |  |  |
| **Data collection** |  |  |  |  |  |  |  |  |
| EM equipment | Titan Krios |  | Titan Krios |  |  | Titan Krios |  |  |
| Voltage (kV) | 300 |  | 300 |  |  | 300 |  |  |
| Detector | Gatan K3 camera |  | Gatan K3 camera |  |  | Gatan K3 camera |  |  |
| Pixel size (Å) | 1.093 |  | 1.093 |  |  | 1.093 |  |  |
| Electron dose (e<sup>-</sup>/Å<sup>2</sup>) | 50.2 |  | 50.2 |  |  | 50.2 |  |  |
| Exposure time (s) | 3 |  | 3 |  |  | 3 |  |  |
| Frames | 30 |  | 30 |  |  | 30 |  |  |
| Defocus range (μm) | -0.8 to -2.5 |  | -0.8 to -2.5 |  |  | -0.8 to -2.5 |  |  |
| **Reconstruction** |  |  |  |  |  |  |  |  |
| Softwares | Relion 3.1& cryoSPARC |  |  |  |  |  |  |  |
| Structures | S-open | S-close | C1 | C2 | RBD-1-ACE2 | S-open-S3H3 | S-close-S3H3 | SD1-S3H3 |
| Final particles | 69,873 | 108,509 | 62,082 | 79,456 | 141,538 | 162,221 | 75,900 | 101,192 |
| Symmetry | C1 | C1 | C1 | C1 | C1 | C1 | C1 | C1 |
| FSC threshold | 0.143 |  |  |  |  |  |  |  |
| Final overall resolution (Å) | 3.21 | 3.08 | 3.69 | 3.66 | 3.67 | 3.48 | 3.64 | 3.61 |
| **Atomic modeling** |  |  |  |  |  |  |  |  |
| Softwares | Rosetta & Phenix & Coot |  |  |  |  |  |  |  |
| Rms deviations |  |  |  |  |  |  |  |  |
| Bond length (Å) | 0.0020 | 0.0020 | 0.0045 | 0.004 | 0.008 | 0.0037 | 0.0036 | 0.0033 |
| Bond Angle (°) | 0.49 | 0.49 | 1.05 | 1.034 | 1.29 | 0.94 | 0.94 | 0.94 |
| Ramachandran plot (%) |  |  |  |  |  |  |  |  |
| Favored | 95.99 | 95.97 | 96.98 | 96.29 | 94.92 | 96.07 | 97.08 | 95.29 |
| Allowed | 3.98 | 4.03 | 3.02 | 3.65 | 4.70 | 3.85 | 2.85 | 4.71 |
| Outliers | 0.00 | 0.00 | 0.00 | 0.05 | 0.38 | 0.08 | 0.00 | 0.00 |
| Molprobity score | 1.43 | 1.46 | 1.40 | 1.74 | 1.66 | 1.70 | 1.67 | 1.72 |
| Clash score | 3.65 | 3.81 | 4.48 | 9.12 | 5.65 | 7.94 | 7.6 | 7.1 |

**Table S2.** Omicron S-close protomer 1 and protomer 3 interactions.

| S-close protomer 1 |  | S-close protomer 3 |  | Interaction | Distance(Å) |
| --- | --- | --- | --- | --- | --- |
| Residue | Atom | Residue | Atom |  |  |
| TYR 396 | [OH] | TYR 200 | [OH] | H-bond | 3.51 |
| ASN 417 | [OD1] | ALA 372 | [N] | H-bond | 2.52 |
| ARG 765 | [NH2] | THR 302 | [O] | H-bond | 3.68 |
| ASN 703 | [N] | ILE 788 | [O] | H-bond | 3.34 |
| ALA 701 | [O] | GLN 787 | [NE2] | H-bond | 2.56 |
| ALA 701 | [O] | ILE 788 | [H] | H-bond | 2.46 |
| ALA 668 | [N] | PRO 863 | [O] | H-bond | 3.80 |
| GLY 669 | [N] | LEU 864 | [O] | H-bond | 3.59 |
| ALA 713 | [N] | GLN 895 | [O] | H-bond | 3.37 |
| SER1123 | [OG] | GLU 918 | [OE2] | H-bond | 3.63 |
| GLU 702 | [OE2] | LYS 790 | [NZ] | H-bond | 2.49 |
| TYR 707 | [OH] | THR 883 | [OG1] | H-bond | 2.81 |
| TYR 707 | [OH] | SER 884 | [OG] | H-bond | 2.65 |
| LYS 547 | [O] | ASN 978 | [ND2] | H-bond | 3.47 |
| LYS 557 | [NZ] | SER 45 | [OG] | H-bond | 3.61 |
| GLN 965 | [NE2] | SER 758 | [OG] | H-bond | 2.44 |
| ARG 408 | [NH2] | PHE 375 | [O] | H-bond | 2.31 |
| ARG 408 | [NH2] | THR 376 | [OG1] | H-bond | 2.57 |
| LYS 969 | [N] | GLN 755 | [O] | H-bond | 3.87 |
| PHE 970 | [N] | GLN 755 | [O] | H-bond | 3.70 |
| ASN 317 | [ND2] | ASP 737 | [OD1] | H-bond | 3.61 |
| ARG 357 | [NH1] | PRO 230 | [O] | H-bond | 3.87 |
| ARG319 | [NH2] | ASP 737 | [OD2] | H-bond | 3.46 |
| LYS 386 | [NZ] | PHE 981 | [O] | H-bond | 2.95 |
| VAL 1040 | [N] | GLU 1031 | [OE2] | H-bond | 3.07 |
| GLU 702 | [OE1] | LYS 790 | [NZ] | Salt bridge | 3.57 |
| GLU 702 | [OE2] | LYS 790 | [NZ] | Salt bridge | 2.49 |
| ARG 319 | [NH1] | ASP 737 | [OD2] | Salt bridge | 3.94 |
| ARG 319 | [NH2] | ASP 737 | [OD2] | Salt bridge | 3.46 |

**Table S3.** Omicron S-close protomer 1 and protomer 2 interactions.

| S-close Protomer 1 |  | S-close Protomer 2 |  | Interaction | Distance(Å) |
| --- | --- | --- | --- | --- | --- |
| Residue | Atom | Residue | Atom |  |  |
| PHE 43 | [O] | ARG 567 | [N] | H-bond | 3.38 |
| TYR 200 | [OH] | TYR 396 | [OH] | H-bond | 3.82 |
| PRO 230 | [O] | ARG 357 | [NH1] | H-bond | 3.74 |
| TYR 369 | [OH] | ASN 460 | [ND2] | H-bond | 3.21 |
| ASN 370 | [OD1] | TYR 421 | [OH] | H-bond | 2.81 |
| PHE 375 | [O] | ARG 408 | [NH1] | H-bond | 3.14 |
| PHE 375 | [O] | ARG 408 | [NH2] | H-bond | 3.18 |
| ASP 737 | [OD2] | ARG 319 | [NH2] | H-bond | 3.78 |
| GLN 755 | [O] | LYS 969 | [N] | H-bond | 3.42 |
| GLN 755 | [O] | SER 968 | [OG] | H-bond | 2.21 |
| GLN 755 | [O] | PHE 970 | [N] | H-bond | 3.26 |
| SER 758 | [OG] | GLN 965 | [NE2] | H-bond | 2.31 |
| ILE 788 | [O] | ASN 703 | [N] | H-bond | 3.35 |
| PRO 863 | [O] | ALA 668 | [N] | H-bond | 3.14 |
| LEU 864 | [O] | ALA 668 | [N] | H-bond | 3.65 |
| LEU 864 | [O] | GLY 669 | [N] | H-bond | 3.07 |
| SER 884 | [OG] | TYR 707 | [OH] | H-bond | 2.77 |
| LEU 894 | [O] | TYR 707 | [OH] | H-bond | 3.53 |
| GLN 895 | [O] | ALA 713 | [N] | H-bond | 3.53 |
| PHE 981 | [O] | LYS 386 | [NZ] | H-bond | 2.40 |
| ARG 983 | [O] | SER 383 | [N] | H-bond | 3.28 |
| GLU 1031 | [OE2] | VAL 1040 | [N] | H-bond | 2.97 |
| LYS 41 | [NZ] | PHE 562 | [O] | H-bond | 3.42 |
| TYR 200 | [OH] | GLU 516 | [OE2] | H-bond | 2.33 |
| TYR 369 | [OH] | ASN 460 | [OD1] | H-bond | 3.52 |
| GLY 757 | [N] | SER 968 | [OG] | H-bond | 3.86 |
| LYS 764 | [NZ] | GLN 314 | [OE1] | H-bond | 2.89 |
| THR 768 | [OG1] | GLN 314 | [OE1] | H-bond | 3.54 |
| GLN 787 | [NE2] | ALA 701 | [O] | H-bond | 2.32 |
| ILE 788 | [N] | ALA 701 | [O] | H-bond | 3.52 |
| LYS 790 | [NZ] | GLU 702 | [OE2] | H-bond | 2.69 |
| LYS 856 | [NZ] | THR 572 | [OG1] | H-bond | 2.80 |
| TYR 904 | [OH] | GLY 1093 | [O] | H-bond | 2.65 |
| SER 975 | [OG] | ASP 571 | [OD2] | H-bond | 2.52 |
| ASN 978 | [ND2] | LYS 547 | [O] | H-bond | 3.08 |
| ASP 737 | [OD2] | ARG 319 | [NH2] | Salt bridge | 3.78 |
| LYS 790 | [NZ] | GLU 702 | [OE1] | Salt bridge | 3.64 |
| LYS 790 | [NZ] | GLU 702 | [OE2] | Salt bridge | 2.69 |

**Table S4.** Omicron RBD-1-ACE2 structure revealed RBD/ACE2 interactions.

| Omicron S RBD-1 |  | ACE2 |  | Interaction | Distance(Å) |
| --- | --- | --- | --- | --- | --- |
| Residue | Atom | Residue | Atom |  |  |
| ASN 487 | [OD1] | TYR 83 | [OH] | H-bond | 2.45 |
| TYR 489 | [OH] | TYR 83 | [OH] | H-bond | 3.00 |
| SER 494 | [O] | HIS 34 | [NE2] | H-bond | 3.09 |
| SER 496 | [O] | LYS 353 | [NZ] | H-bond | 3.65 |
| SER 496 | [OG] | LYS 353 | [NZ] | H-bond | 2.82 |
| ASN 477 | [ND2] | SER 19 | [O] | H-bond | 3.86 |
| ASN 477 | [ND2] | SER 19 | [OG] | H-bond | 3.38 |
| ASN 487 | [ND2] | GLN 24 | [OE1] | H-bond | 2.74 |
| ARG 493 | [NH2] | GLU 35 | [OE1] | H-bond | 3.58 |
| TYR 449 | [OH] | ASP 38 | [OD1] | H-bond | 2.79 |
| ARG 498 | [NH1] | ASP 38 | [OD1] | H-bond | 3.37 |
| TYR 449 | [OH] | ASP 38 | [OD2] | H-bond | 2.62 |
| TYR 449 | [OH] | GLN 42 | [OE1] | H-bond | 3.27 |
| ARG 498 | [NH1] | GLN 42 | [OE1] | H-bond | 3.04 |
| HIS 505 | [ND1] | LYS 353 | [O] | H-bond | 2.85 |
| THR 500 | [OG1] | ASP 355 | [OD2] | H-bond | 3.40 |
| ARG 493 | [NH2] | GLU 35 | [OE1] | Salt bridge | 3.58 |
| ARG 493 | [NH2] | GLU 35 | [OE2] | Salt bridge | 3.93 |
| ARG 498 | [NH1] | ASP 38 | [OD1] | Salt bridge | 3.37 |

**Table S5.** Contacting residues (a sidechain distance cut off 4 Å) at the Omicron RBD/ACE2 interface.

| Omicron S RBD-1 | ACE2 |
| --- | --- |
| Y449 | D38, Q42 |
| Y453 | H34 |
| L455 | D30 |
| F456 | T27, D30, K31 |
| A475 | Q24, T27 |
| N477 | S19 |
| F486 | M82, Y83 |
| N487 | Q24, Y83 |
| Y489 | T27, F28 |
| R493 | H34, E35 |
| S494 | H34 |
| S496 | D38, K353 |
| R498 | D38, Y41, Q42 |
| T500 | Y41, D355, R357 |
| Y501 | Y41, K353, G354, D355 |
| G502 | G354 |
| H505 | K353, G354 |

**Table S6.** Omicron S SD1-S3H3 structure revealed S/S3H3 interactions.

| Omicron S |  | S3H3 |  | Interaction | Distance(Å) |
| --- | --- | --- | --- | --- | --- |
| Residue | Atom | Residue | Atom |  |  |
| LYS 537 | [NZ] | ASP 55 | [OD2] | H-bond | 3.35 |
| THR 581 | [OG1] | ASP 102 | [OD1] | H-bond | 3.25 |
| SER 555 | [O] | SER 32 | [OG] | H-bond | 3.05 |
| ILE 584 | [O] | SER 32 | [OG] | H-bond | 3.80 |
| LYS 537 | [NZ] | ASP 55 | [OD2] | Salt bridge | 3.35 |

Heavy chain
Light chain

**Table S7.** Contacting residues (a sidechain distance cut off 4 Å) at the Omicron SD1/S3H3 interface.

| Omicron S | S3H3 |
| --- | --- |
| T323 | S54 |
| E324 | R31 |
| N532 | F32 |
| L533 | Y101 |
| V534 | R31, W33 |
| K535 | Y101 |
| N536 | W33, R59, L98 |
| K537 | H52, D55 |
| E554 | Y36, S95, R96 |
| S555 | A31, S32 |
| N556 | A31, R96 |
| T581 | D102 |
| L582 | Y34 |
| E583 | Y103 |
| I584 | S32 |

Heavy chain
Light chain

## References

1 Grabowski, F., Preibisch, G., Gizinski, S., Kochanczyk, M. & Lipniacki, T. SARS-CoV-2 Variant of Concern 202012/01 Has about Twofold Replicative Advantage and Acquires Concerning Mutations. Viruses 13, doi:10.3390/v13030392 (2021).

2 Wise, J. Covid-19: New coronavirus variant is identified in UK. Bmj 371, m4857, doi:10.1136/bmj.m4857 (2020).

3 Davies, N. G. et al. Estimated transmissibility and impact of SARS-CoV-2 lineage B.1.1.7 in England. Science 372, doi:10.1126/science.abg3055 (2021).

4 Gobeil, S. M. et al. Effect of natural mutations of SARS-CoV-2 on spike structure, conformation, and antigenicity. Science, doi:10.1126/science.abi6226 (2021).

5 Cai, Y. et al. Structural basis for enhanced infectivity and immune evasion of SARS-CoV-2 variants. Science, doi:10.1126/science.abi9745 (2021).

6 Yuan, M. et al. Structural and functional ramifications of antigenic drift in recent SARS-CoV-2 variants. Science, doi:10.1126/science.abh1139 (2021).

7 Tegally, H. et al. Detection of a SARS-CoV-2 variant of concern in South Africa. Nature 592, 438–443, doi:10.1038/s41586-021-03402-9 (2021).

8 Msomi, N., Mlisana, K., de Oliveira, T. & Network for Genomic Surveillance in South Africa writing, g. A genomics network established to respond rapidly to public health threats in South Africa. Lancet Microbe 1, e229–e230, doi:10.1016/S2666-5247(20)30116-6 (2020).

9 Voloch, C. M. et al. Genomic characterization of a novel SARS-CoV-2 lineage from Rio de Janeiro, Brazil. Journal of virology, doi:10.1128/JVI.00119-21 (2021).

10 Singh, J., Rahman, S. A., Ehtesham, N. Z., Hira, S. & Hasnain, S. E. SARS-CoV-2 variants of concern are emerging in India. Nat Med 27, 1131–1133, doi:10.1038/s41591-021-01397-4 (2021).

11 Winger, A. & Caspari, T. The Spike of Concern-The Novel Variants of SARS-CoV-2. Viruses 13, doi:10.3390/v13061002 (2021).

12 Grabowski, F., Kochańczyk, M. & Lipniacki, T. Omicron strain spreads with the doubling time of 3.2—3.6 days in South Africa province of Gauteng that achieved herd immunity to Delta variant. medRxiv, doi:10.1101/2021.12.08.21267494 (2021).

13 Burki, T. K. Omicron variant and booster COVID-19 vaccines. The Lancet Respiratory Medicine, doi:10.1016/s2213-2600(21)00559-2 (2021).

14 WHO. Enhancing readiness for Omicron (B. 1.1.529): Technical brief and priority actions for Member States. https://www.who.int/publications/m/item/enhancing-readiness-for-omicron-(b.1.1.529)-technical-brief-and-priority-actions-for-member-states (2021).

15 Cameroni, E. et al. Broadly neutralizing antibodies overcome SARS-CoV-2 Omicron antigenic shift. Nature, doi:10.1038/d41586-021-03825-4 (2021).

16 Mannar, D. et al. SARS-CoV-2 Omicron Variant: ACE2 Binding, Cryo-EM Structure of Spike Protein-ACE2 Complex and Antibody Evasion. BioRxiv, doi:10.1101/2021.12.19.473380 (2021).

17 Hoffmann, M. et al. The Omicron variant is highly resistant against antibody-mediated neutralization – implications for control of the COVID-19 pandemic. Cell, doi:10.1016/j.cell.2021.12.032 (2021).

18 Carreño, J. M. et al. Activity of convalescent and vaccine serum against SARS-CoV-2 Omicron. Nature, doi:10.1038/d41586-021-03846-z (2021).

19 Liu, L. et al. Striking antibody evasion manifested by the Omicron variant of SARS-CoV-2. Nature, doi:10.1038/d41586-021-03826-3 (2021).

20 Cao, Y. et al. Omicron escapes the majority of existing SARS-CoV-2 neutralizing antibodies. Nature, doi:10.1038/d41586-021-03796-6 (2021).

21 Planas, D. et al. Considerable escape of SARS-CoV-2 Omicron to antibody neutralization. Nature, doi:10.1038/d41586-021-03827-2 (2021).

22 Cele, S. et al. Omicron extensively but incompletely escapes Pfizer BNT162b2 neutralization. Nature, doi:10.1038/d41586-021-03824-5 (2021).

23 Tang, T., Bidon, M., Jaimes, J. A., Whittaker, G. R. & Daniel, S. Coronavirus membrane fusion mechanism offers a potential target for antiviral development. Antiviral Res 178, 104792, doi:10.1016/j.antiviral.2020.104792 (2020).

24 Rabaan, A. A. et al. SARS-CoV-2, SARS-CoV, and MERS-COV: A comparative overview. Infez Med 28, 174–184 (2020).

25 Wang, Q. et al. Structural and Functional Basis of SARS-CoV-2 Entry by Using Human ACE2. Cell 181, 894–904 e899, doi:10.1016/j.cell.2020.03.045 (2020).

26 Lan, J. et al. Structure of the SARS-CoV-2 spike receptor-binding domain bound to the ACE2 receptor. Nature 581, 215–220, doi:10.1038/s41586-020-2180-5 (2020).

27 Shang, J. et al. Structural basis of receptor recognition by SARS-CoV-2. Nature 581, 221–224, doi:10.1038/s41586-020-2179-y (2020).

28 Yin, W. et al. Structures of the Omicron Spike trimer with ACE2 and an anti-Omicron antibody. bioRxiv, 2021.2012.2027.474273, doi:10.1101/2021.12.27.474273 (2021).

29 Xu, S. et al. Mapping cross-variant neutralizing sites on the SARS-CoV-2 spike protein. Emerging Microbes & Infections, 1–51, doi:10.1080/22221751.2021.2024455 (2021).

30 Zhang, J. et al. Structural impact on SARS-CoV-2 spike protein by D614G substitution. Science 372, 525–530, doi:10.1126/science.abf2303 (2021).

31 Wang, Y. et al. Conformational dynamics of the Beta and Kappa SARS-CoV-2 spike proteins and their complexes with ACE2 receptor revealed by cryo-EM. Nat Commun doi:10.1038/s41467-021-27350-0 (2021).

32 Wang, Y. et al. Structural basis for SARS-CoV-2 Delta variant recognition of ACE2 receptor and broadly neutralizing antibodies. Nat Commun (Accepted in principle) (2022).

33 Toelzer, C. et al. Free fatty acid binding pocket in the locked structure of SARS-CoV-2 spike protein. Science 370, 725–730, doi:10.1126/science.abd3255 (2020).

34 Wrobel, A. G. et al. SARS-CoV-2 and bat RaTG13 spike glycoprotein structures inform on virus evolution and furin-cleavage effects. Nature structural & molecular biology 27, 763–767, doi:10.1038/s41594-020-0468-7 (2020).

35 Cai, Y. et al. Distinct conformational states of SARS-CoV-2 spike protein. Science, doi:10.1126/science.abd4251 (2020).

36 Xu, C. et al. Conformational dynamics of SARS-CoV-2 trimeric spike glycoprotein in complex with receptor ACE2 revealed by cryo-EM. Sci Adv 7, doi:10.1126/sciadv.abe5575 (2021).

37 Punjani, A., Rubinstein, J. L., Fleet, D. J. & Brubaker, M. A. cryoSPARC: algorithms for rapid unsupervised cryo-EM structure determination. Nat Methods 14, 290–296, doi:10.1038/nmeth.4169 (2017).

38 Cui, Z. et al. Structural and functional characterizations of altered infectivity and immune evasion of SARS-CoV-2 Omicron variant. BioRxiv, doi: 10.1101/2021.12.29.474402 (2021).

39 McCallum, M. et al. Structural basis of SARS-CoV-2 Omicron immune evasion and receptor engagement. BioRxiv, doi:10.1101/2021.12.28.474380 (2021).

40 Lan, J. et al. Structural and computational insights into the SARS-CoV-2 Omicron RBD-ACE2 interaction. BioRxiv, doi:10.1101/2022.01.03.474855 (2022).

41 Han, P. et al. Receptor binding and complex structures of human ACE2 to spike RBD from Omicron and Delta SARS-CoV-2. Cell, doi:10.1016/j.cell.2022.01.001 (2022).

42 Mannar, D. et al. Structural analysis of receptor binding domain mutations in SARS-CoV-2 variants of concern that modulate ACE2 and antibody binding. Cell Rep 37, 110156, doi:10.1016/j.celrep.2021.110156 (2021).

43 Laffeber, C., de Koning, K., Kanaar, R. & Lebbink, J. H. G. Experimental Evidence for Enhanced Receptor Binding by Rapidly Spreading SARS-CoV-2 Variants. Journal of molecular biology 433, 167058, doi:10.1016/j.jmb.2021.167058 (2021).

44 Zhang, C. et al. Development and structural basis of a two-MAb cocktail for treating SARS-CoV-2 infections. Nature Communications 12, doi:10.1038/s41467-020-20465-w (2021).

45 Zeng, C. et al. Neutralization and Stability of SARS-CoV-2 Omicron Variant. BioRxiv, doi: 10.1101/2021.12.16.472934 (2021).

46 Meng, B. et al. SARS-CoV-2 Omicron spike mediated immune escape, infectivity and cell-cell fusion. BioRxiv, doi: 10.1101/2021.12.17.473248 (2021).

47 Kwong, P. D. et al. HIV-1 evades antibody-mediated neutralization through conformational masking of receptor-binding sites. Nature 420, 678–682, doi:10.1038/nature01188 (2002).

48 Munro, J. B. et al. Conformational dynamics of single HIV-1 envelope trimers on the surface of native virions. Science 346, 759–763, doi:10.1126/science.1254426 (2014).

49 Barnes, C. O. et al. SARS-CoV-2 neutralizing antibody structures inform therapeutic strategies. Nature 588, 682–687, doi:10.1038/s41586-020-2852-1 (2020).

50 Joshi, N., Tyagi, A. & Nigam, S. Molecular Level Dissection of Critical Spike Mutations in SARS-CoV-2 Variants of Concern (VOCs): A Simplified Review. ChemistrySelect 6, 7981–7998, doi:10.1002/slct.202102074 (2021).

51 Campbell, F. et al. Increased transmissibility and global spread of SARS-CoV-2 variants of concern as at June 2021. Euro Surveill 26, doi: 10.2807/1560-7917.ES.2021.26.24.2100509 (2021).

52 Zheng, S. Q. et al. MotionCor2: anisotropic correction of beam-induced motion for improved cryo-electron microscopy. Nat Methods 14, 331–332, doi:10.1038/nmeth.4193 (2017).

53 Fernandez-Leiro, R. & Scheres, S. H. W. A pipeline approach to single-particle processing in RELION. Acta Crystallogr D Struct Biol 73, 496–502, doi:10.1107/S2059798316019276 (2017).

54 Rohou, A. & Grigorieff, N. CTFFIND4: Fast and accurate defocus estimation from electron micrographs. J Struct Biol 192, 216–221, doi:10.1016/j.jsb.2015.08.008 (2015).

55 Emsley, P. & Cowtan, K. Coot: model-building tools for molecular graphics. Acta Crystallogr D Biol Crystallogr 60, 2126–2132, doi:10.1107/S0907444904019158 (2004).

56 DiMaio, F. et al. Atomic-accuracy models from 4.5-A cryo-electron microscopy data with density-guided iterative local refinement. Nature methods 12, 361–365, doi:10.1038/nmeth.3286 (2015).

57 Adams, P. D. et al. PHENIX: a comprehensive Python-based system for macromolecular structure solution. Acta Crystallogr D Biol Crystallogr 66, 213–221, doi:10.1107/S0907444909052925 (2010).

58 Krissinel, E. & Henrick, K. Inference of macromolecular assemblies from crystalline state. Journal of molecular biology 372, 774–797, doi:10.1016/j.jmb.2007.05.022 (2007).

59 Pettersen, E. F. et al. UCSF Chimera--a visualization system for exploratory research and analysis. J Comput Chem 25, 1605–1612, doi:10.1002/jcc.20084 (2004).

60 Goddard, T. D. et al. UCSF ChimeraX: Meeting modern challenges in visualization and analysis. Protein Sci 27, 14–25, doi:10.1002/pro.3235 (2018).

